# Parkin-mediated ubiquitination contributes to the constitutive turnover of mitochondrial fission factor (Mff)

**DOI:** 10.1101/553859

**Authors:** Laura Lee, Richard Seager, Yasuko Nakamura, Kevin A. Wilkinson, Jeremy M. Henley

## Abstract

The mitochondrial outer membrane protein Mitochondrial Fission Factor (Mff) plays a key role in both physiological and pathological fission. It is well established that in stressed or functionally impaired mitochondria the PINK1 recruits the ubiquitin ligase Parkin which ubiquitinates Mff to facilitate the removal of defective mitochondria and maintain the integrity mitochondrial network. Here we show that, in addition to this clearance pathway, Parkin also ubiquitinates Mff in a PINK1-dependent manner under basal, non-stressed conditions to regulate constitutive Mff turnover. We further show that removing Parkin with shRNA knockdown does not completely prevent Mff ubiquitination under these conditions indicating that at least one other ubiquitin ligase contributes to Mff proteostasis. These data demonstrate that Parkin plays a role in physiological maintenance of mitochondrial membrane protein composition in healthy mitochondria through constitutive low-level activation.

## Introduction

Mitochondria are double membrane-bound organelles that generate 90% of cellular ATP (1). In most cells, mitochondria form extensive and dynamic networks, undergoing continuous cycles of fission and fusion. This creates a highly adaptable and efficient energy transfer system to rapidly deliver ATP to where it is most needed (2). In addition, fission plays a central role in the sequestration and selective degradation of defective mitochondria by mitophagy (3, 4). Mitochondrial fission and fusion are both tightly regulated processes. Mitochondrial dynamics is largely orchestrated by the GTPases. Dynamin and dynamin-related protein (Drp) mediate fission (5, 6) whereas fusion of the mitochondrial outer and inner membranes is driven by the GTPases mitofusins (Mfn) 1 and 2 and Opa1 respectively (7).

Dynamin-related protein 1 (Drp1) is predominantly a cytosolic protein with only ∼3% bound to mitochondria under basal conditions (8). Nonetheless, in cells lacking functional Drp1 the equilibrium between fission and fusion is perturbed leading to highly elongated and interconnected mitochondria, largely localised in perinuclear clusters (9). During cell stress, rates of fission and fragmentation increase leading to the release of pro-apoptotic cytochrome *c* from mitochondria, a process that can be delayed by mutation or deletion of Drp1 (10).

Because Drp1 lacks the membrane targeting PH-domain present in conventional dynamins it requires membrane-bound adaptor/receptor proteins to be recruited to the mitochondrial outer membrane (MOM) (11). Four mitochondrial Drp1 receptors have been identified; Fis1, MiD49, MiD51 and Mff (12). Of these, Fis1 is dispensable for mammalian mitochondrial fission (13). The MiD proteins are specific to higher eukaryotes and although they can each recruit Drp1 to mitochondrial fission sites (14, 15) it remains unclear if MiD proteins facilitate fusion or inhibit fission (16). Mff facilitates the majority of Drp1 recruitment and is the best characterised Drp1 receptor. It is a ∼35kDa protein with a single C-terminal transmembrane domain and interacts with Drp1 via its N-terminus (17). Like Drp1-null cells, Mff-knockout cells have grossly elongated mitochondria under basal conditions, and attenuated fragmentation and apoptosis following stress (18).

Parkin is a ubiquitin ligase that is inactive in the cytosol but is recruited to damaged/depolarised mitochondria where it is activated by the MOM protein PTEN-induced protein kinase 1 (PINK1). PINK1 is basally maintained at very low levels by rapid proteolytic degradation soon after mitochondrial import (19, 20). However, loss of membrane potential in damaged or defective mitochondrial inhibits PINK1-proteolysis, resulting in its accumulation on the outer membrane, where it phosphorylates mitochondrial u ubiquitin at Serine 65 and triggers mitophagy (21, 22).

Briefly, PINK1-phosphorylated ubiquitin (pUb) binds to and alters the conformation of Parkin. This makes Serine 65 within the Ubiquitin-like domain (UbL) accessible for PINK1-mediated phosphorylation, which initiate a cascade of subsequent conformational changes exposing the catalytic site of Parkin (22–24). In a positive-feedback loop, Parkin ubiquitinates mitochondrial proteins, providing further substrates for PINK1-mediated phosphorylation, which then recruit more Parkin (25, 26). For example, mitophagy induced by the mitochondrial proton gradient uncoupler carbonyl cyanide m-chlorophenyl hydrazine (CCCP) is largely dependent on Parkin-mediated, non-selective ubiquitination of mitochondrial proteins with K48- and K63-linked ubiquitin chains (27, 28). Mitochondrial depolarisation leads to PINK1 accumulation on the surface of mitochondria that recruits Parkin to indiscriminately tags MOM proteins with K48-linked ubiquitin chains, marking them for excision and proteasomal degradation (27, 29). The remaining portion of the mitochondrion is then tagged with K63-linked ubiquitin that recruits phagosomal adaptors including p62 (30) resulting in the engulfment of the organelle into an autophagosome prior to lysosomal fusion and degradation (31, 32). Thus, this elegant quality control mechanism identifies damaged mitochondria and targets proteins for degradation.

Moreover, in cells lacking functional PINK1 and/or Parkin mitochondria undergo fragmentation due to excessive Drp1-mediated fission (33–35). However, the roles of Parkin in non-stressed mitochondria has not been extensively investigated. Here we show that, independent of stress-induced mitophagy, Mff is ubiquitinated by Parkin and at least one other E3 ligase under basal conditions. Our data indicate that Parkin-mediated ubiquitination triggers lysosomal degradation of Mff suggesting a role for Parkin in homeostatic maintenance of Mff levels and mitochondrial integrity.

## Materials and Methods

### Molecular Biology

21bp short hairpin (shRNA) constructs used in Figs 1–4 (targeting human shParkin: 5’- ACCAGCATCTTCCAGCTCAAG-3’; non-specific shControl: 5’- AACGTACGCGGAATACTTCGA-3’) were cloned under a H1 promoter into a modified pSUPER vector co-expressing mCherry driven by an SFFV promoter. Alternative Parkin shRNAs (SI Fig. 1b) were cloned in the same way, with target sequences 5’- GCTTAGACTGTTTCCACTTAT-3’ (Parkin (Berger)) and 5’-AACTCCAGCCATGGTTTCCCA-3’ (Parkin (other)). Parkin (Berger) target sequence was previously published (36). Other Parkin shRNA target sequences were designed as part of this study. PINK1 knockdown was performed using MISSION® esiRNA human PINK1 (EHU057101, Sigma Aldrich). Mff knockdown (SI Fig. 1a) was performed using siRNA with the target sequence 5’-CCAUUGAAGGAACGUCAGA-3’ (Eurofins genomics). Firefly Luciferase siRNA was used as a negative control (MISSION® esiRNA Firefly Luciferase, EHUFLUC, Sigma Aldrich).

**Fig. 1.**
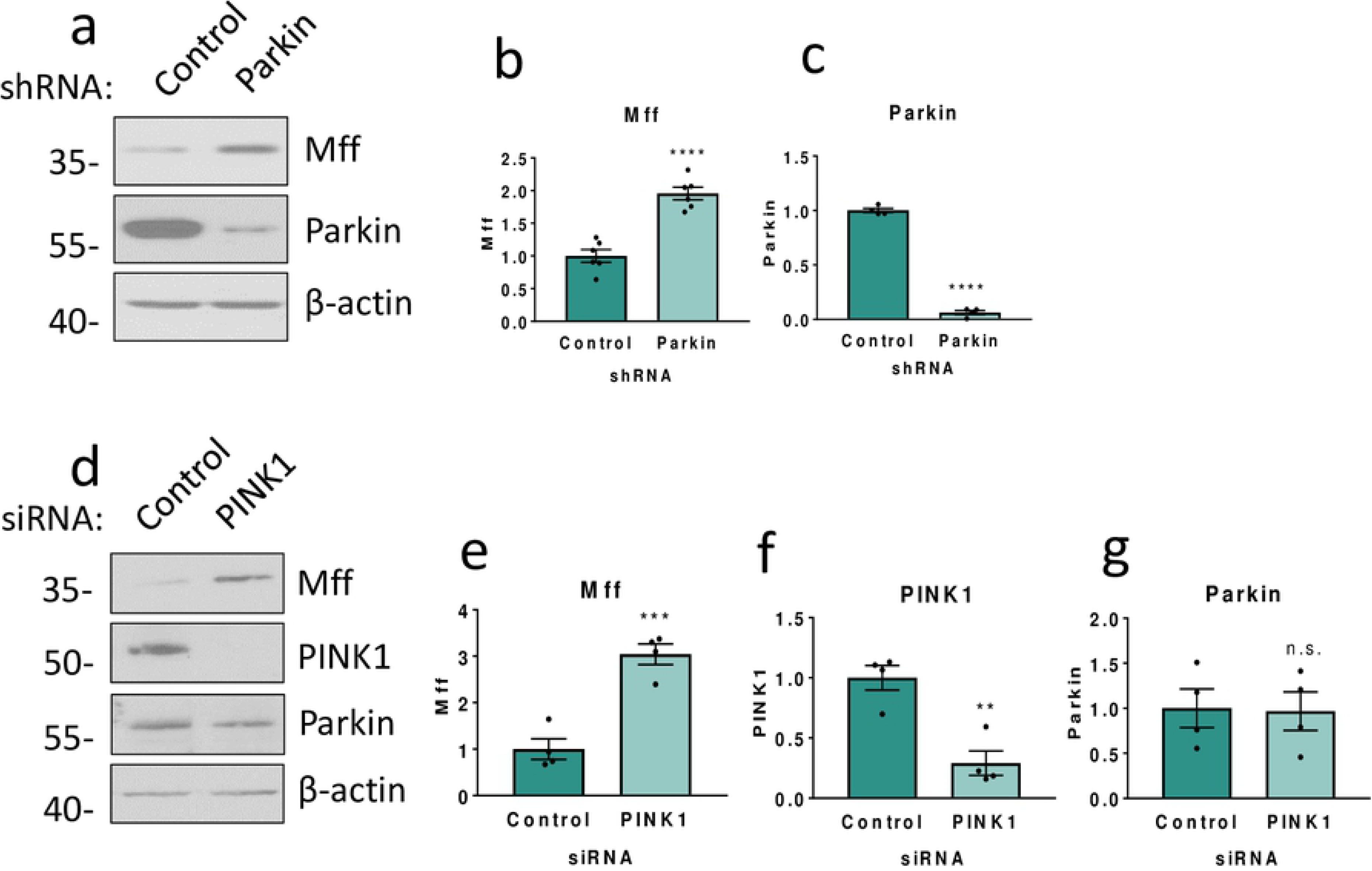
The PINK1/Parkin pathway is involved in Mff stability. a) HEK293T cells were transfected with either shRNA targeting human Parkin or a scrambled control shRNA. Western blots for Mff, Parkin and β-actin. N= 4-7. Quantitative analysis of Mff (b) and Parkin (c) levels using Student’s unpaired t-test. d) HEK293T cells were transfected with either siRNA targeting human PINK1 or a control siRNA (firefly luciferase). Western blots for Mff, PINK1, Parkin and β-actin. N=4. Quantitative analysis of Mff (e), PINK1 (f) and Parkin (g). Data presented as mean ± SEM. ** p < 0.01, *** p < 0.001, **** p < 0.0001.

The open reading frame of human Mff (isoform I, accession number: Q9GZY8) was cloned into pECFP between 5’ KpnI and 3’ BamHI restriction sites. CFP-Mff expression was driven by a CMV promoter. The open reading frame of human Parkin (full length, accession number: O60260) was cloned into pcDNA3.1(+) between 5’ HindIII and 3’ BamHI restriction sites. Parkin expression was driven by a CMV promoter. CFP-Mff K151R, K302R and 2KR, Parkin S65A and S65D were generated by site-directed mutagenesis.

### HEK293T cell culture and transfection

Human Embryonic Kidney (HEK293T) cells were obtained from The European Collection of Cell Cultures (ECACC). Cultures were maintained at 37°C in a humidified cell culture incubator, supplied with 5% CO_2_ and 95% O_2_, in Dulbecco’s Modified Eagle’s Medium (Lonza) supplemented with 10% (v/v) Foetal Bovine Serum (Sigma) and 2mM L-Glutamine (Gibco). For transfection, cells were plated on dishes pre-coated with 0.1mg/mL poly-L-lysine to promote adhesion. Lipofectamine 2000 transfection reagent (Invitrogen) was used according to manufacturer’s protocol. Cells were lysed 48 hours (protein over-expression) or 72 hours (protein knockdown) post-transfection.

### Protein biochemistry

For immunoblotting, cells were lysed in sample buffer (1x concentrate) containing 2% SDS (w/v), 5% glycerol (v/v), 62.5mM Tris-HCl pH6.8 and 5% (v/v) 2-β-mercaptoethanol. Lysates were heated to 95°C for 10 minutes prior to gel electrophoresis. For immunoprecipitation, cells were lysed in lysis buffer containing 20mM Tris pH7.4, 137mM NaCl, 2mM sodium pyrophosphate, 2mM EDTA, 1% (v/v) Triton X-100, 0.1% (w/v) SDS, 25mM β-glycerophosphate, 10% glycerol (v/v), 1x cOmplete™ protease inhibitor cocktail (Roche) and 20mM N- Ethylmaleimide (NEM, Sigma). Lysates were incubated with GFP-Trap® agarose beads (ChromoTek) at 4°C for 90 minutes with gentle agitation. Beads were pelleted, washed 3 times with wash buffer (lysis buffer without protease inhibitor cocktail or NEM) and unbound material aspirated. 2x concentrate sample buffer was used to elute immunoprecipitated proteins from the beads. Samples were heated to 95°C for 10 minutes prior to gel electrophoresis.

Denaturing SDS-PAGE was performed on 10% (v/v) poly-acrylamide gels, Western blotted PDVF membranes were blocked in 5% (w/v) non-fat milk powder or 4% (w/v) Bovine Serum Albumin (BSA, Sigma) in PBS-T. Primary antibodies used were: Parkin (mouse monoclonal, 1:1000, Santa Cruz sc-32282), Mff (mouse monoclonal, 1:1000, Santa Cruz sc-398731), PINK1 (rabbit monoclonal, 1:1000, Cell Signalling D83G 6946), ubiquitin (mouse monoclonal, 1:1000, P4D1 3936S), GFP (rat monoclonal, 1:10,000, Chromotek 3H9), β-actin (mouse monoclonal, 1:10,000, Sigma-Aldrich A5441). For protein detection, membranes were incubated with HRP-conjugated secondary antibodies (1:10,000; Sigma-Aldrich) and visualised by enhanced chemiluminescence. Protein bands were quantified by densitometry using ImageJ (NIH). Mff- and Parkin-antibody specificity was validated by endogenous protein knockdown (SI Fig. 1).

## Results

### Knockdown of PINK1 or Parkin increases expression of Mff

We first investigated the effects of Parkin knockdown on Mff by transfecting HEK293T cells with plasmids encoding an shRNA sequence targeted to the human Parkin transcript (shParkin) or a control shRNA sequence. shParkin reduced Parkin levels to 6% of control levels 72 hours after transfection (Fig. 1a,c). Consistent with Parkin-mediated ubiquitination and degradation of Mff there was a corresponding increase in levels of Mff (Fig. 1a,b). To exclude the possibility of the Mff increase being due off-target effects of the Parkin shRNA, two further Parkin shRNAs were generated and tested in HEK293T cells, with the same effect on Mff levels (SI Fig. 1b).

To determine if the effect of Parkin on Mff levels was PINK1-dependent we next knocked down PINK1 in HEK293T cells. 72 hours post-transfection, PINK1 was significantly knocked down to 30% of control values (Fig. 1d, f). As expected, loss of PINK1 also significantly increased total levels of Mff (Fig. 1e). Moreover, total levels of Parkin were unaffected by knockdown of PINK1, indicating that it is Parkin activity, rather than expression, that causes the increase in Mff (Fig. 1g).

### Knockdown of Parkin reduces steady-state ubiquitination of Mff

Parkin has been reported to ubiquitinate Mff in HEK293 cells over-expressing HA-tagged Parkin and treated with CCCP to induce mitochondrial depolarisation (30). Our data suggest that the PINK1/Parkin pathway may also control Mff levels in the absence of global cell stress. To confirm if Parkin regulates Mff expression under basal, non-stressed conditions HEK293T cells were co-transfected with CFP-tagged Mff and shRNA targeting Parkin or a scrambled control. 72 hours after transfection, cells were lysed and CFP-Mff retained on GFP-Trap® agarose beads. Steady-state levels of Mff ubiquitination were significantly decreased by Parkin knockdown (Fig. 2a,b). These data demonstrate a role for Parkin in Mff ubiquitination that is not dependent on global mitochondrial stress.

**Fig. 2.**
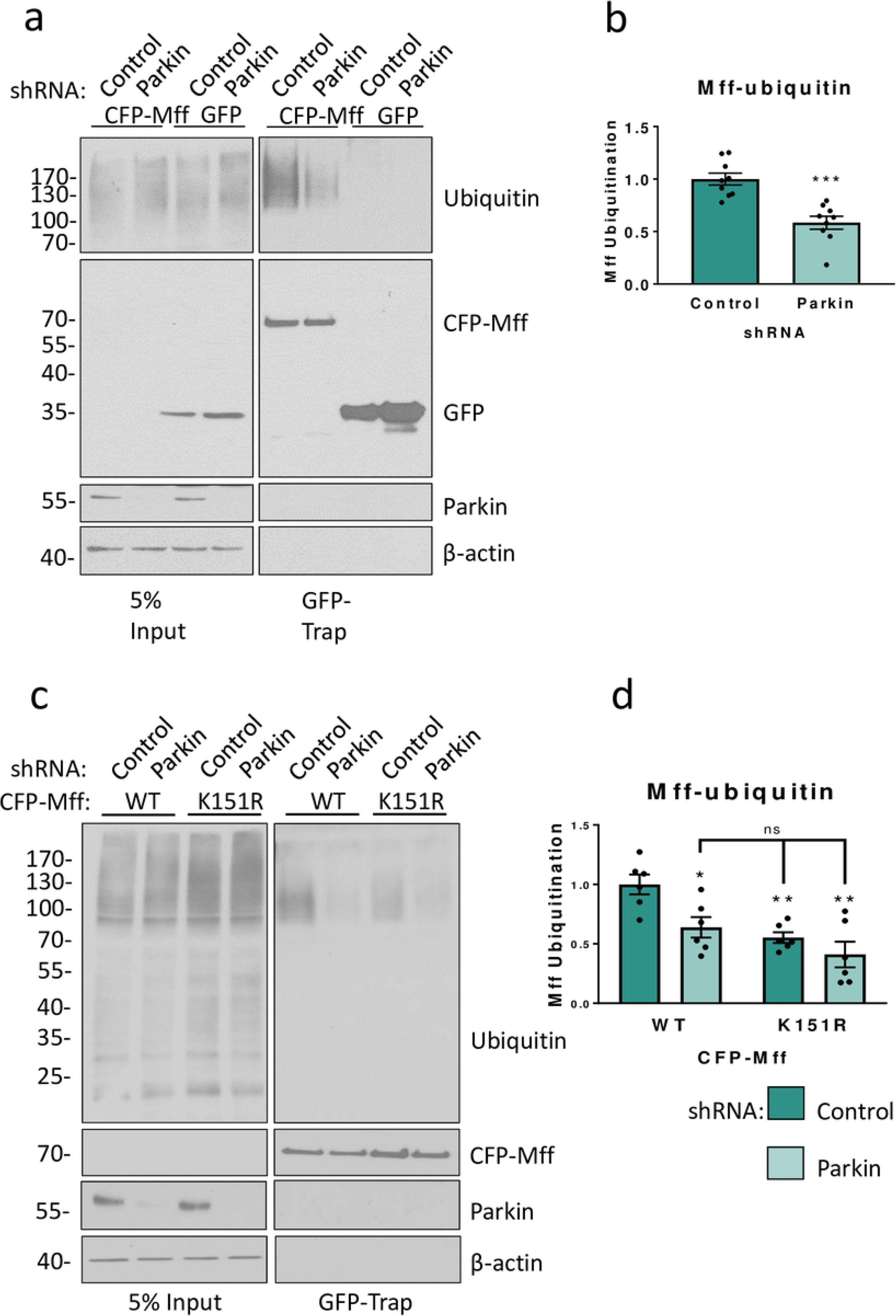
Parkin ubiquitinates Mff at K151 under basal conditions. a) HEK293T cells were co-transfected with CFP-Mff/GFP and shRNA (scrambled control or Parkin-targeting). Western blots of GFP-immunoprecipitation. CFP-tagged Mff immunoprecipitates with endogenous, covalently attached ubiquitin. b) Knockdown of Parkin significantly reduces Mff ubiquitination. N=9. Analysed using unpaired two-tailed students’ t-test. Data presented as mean ± SEM. p < 0.001. c) GFP-immunoprecipitations of exogenously expressed CFP-Mff WT or K151R in HEK293T cells reveal that the K151R mutant has significantly reduced endogenous ubiquitination compared to the WT. d) Quantitative analysis showing that knockdown of Parkin significantly reduces ubiquitination of WT CFP-Mff, but not CFP-Mff K151R. N=6. Analysed using ordinary two-way ANOVA with Tukey’s correction for multiple comparisons with a pooled variance. * p < 0.05, ** p ≤ 0.01.

### Parkin ubiquitinates Mff at K151

The report showing Mff is ubiquitinated by over-expressed Parkin and CCCP-treatment identified K251 as the site of ubiquitination (30). The Mff isoform used in that study (Isoform II, 291 amino acids) is a truncated protein, lacking a large portion of the N-terminus as well as a central region. Here we used Mff isoform I, which is the full-length protein comprising 342 amino acids. Accounting for these differences in the length of the isoforms, K251 in Isoform II corresponds to K302 in Isoform I (SI Fig. 2).

We therefore generated a CFP-Mff mutant in which K302 is replaced with a non-modifiable arginine (K302R). HEK293T cells were co-transfected with CFP-Mff (WT or K302R). 48 hours post-transfection, cells were lysed and GFP-Trap® agarose beads used to precipitate CFP-Mff. Mutation of Mff K302 to arginine (K302R) reduced, but did not abolish, ubiquitination of Mff isoform I (SI Fig. 3).

**Fig. 3.**
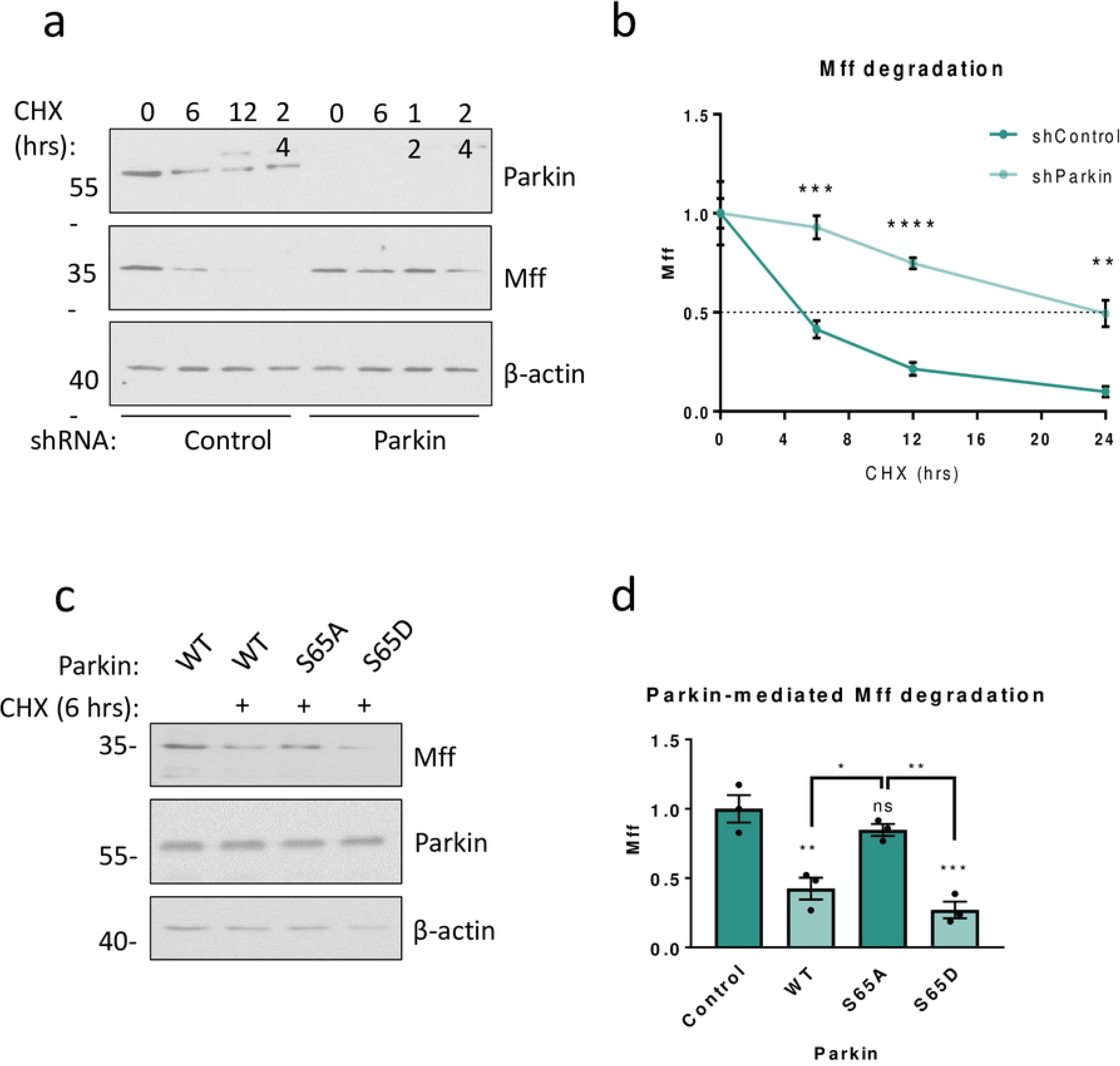
Parkin mediates PINK1-dependent degradation of Mff. a) HEK293T cells were transfected with either scrambled shRNA (control) or shRNA targeting human Parkin. Prior to lysis 72 hours post-transfection, cells were treated with 25µg/mL cycloheximide (CHX) for 0, 6, 12 or 24 hours (0-hour CHX received 24-hour DMSO treatment). Lysates were then Western blotted for Parkin, Mff and β-actin. b) Quantitative analysis of (a), data presented as mean ± SEM. Analysed using unpaired two-tailed Student’s t-tests. N=4. ** p < 0.01, *** p < 0.001, **** p < 0.0001. c) HEK293T cells were transfected with WT or mutant Parkin. Prior to lysis 48 hours post-transfection, cells were treated with 25µg/mL cycloheximide (CHX) for 6 hours (control treated with DMSO, 6 hours). Lysates were then Western blotted for Mff, Parkin and β-actin. d) Quantitative analysis of (c), data presented as mean ± SEM. Analysed using one-way ANOVA with Tukey’s correction for multiple comparisons with a pooled variance. N=3. * p < 0.05, ** p < 0.01, *** p < 0.001.

A recent study has reported that Mff is phosphorylated by AMPK at conserved sites S155 and S172 (highlighted in SI Fig. 2) (37). We hypothesised that, given its conservation and density of modifiable residues, this region could be a ‘hotspot’ for Mff post-translational modifications. We therefore selected K151 as an alternative likely target for ubiquitination. Replacement of lysine 151 with arginine (K151R) also reduced, but failed to abolish, Mff-ubiquitination, as did a double mutant (2KR = K151R and K302R) (SI Fig. 3). These results showing that dual mutation of K151 and K302 both reduce but do not prevent ubiquitination indicate that Mff isoform I has several sites of ubiquitination, K151 and K302 plus at least one other lysine.

To specifically test if Parkin ubiquitinates Mff at K151 under basal conditions, HEK293T cells were co-transfected with CFP-Mff (WT or K151R) and either Parkin or control shRNA. 72 hours post-transfection, cells were lysed and GFP-Trap® agarose beads used to precipitate CFP-Mff. Parkin knockdown significantly reduced ubiquitination of WT CFP-Mff (Fig. 2a,b). However, Parkin knockdown had no significant effect on ubiquitination of CFP-Mff K151R, indicating that under these conditions Parkin preferentially ubiquitinates Mff at K151 (Fig. 2c,d). Interestingly, while, CFP-Mff K151R had significantly reduced ubiquitination compared to WT, some ubiquitin modification was still present. This finding supports data presented in SI Fig. 3 indicating multiple ubiquitination sites and that at least one other ubiquitin ligase can ubiquitinate CFP-Mff at a different lysine from the Parkin targeted K151.

### Parkin mediates PINK1-dependent turnover of Mff

Our results show that Parkin knockdown of significantly decreases ubiquitination and increases levels of Mff levels. We next directly tested the role of Parkin in Mff degradation. HEK293T cells were transfected with Parkin-targeted shRNA or a scrambled shRNA control. Prior to lysis 72 hours post-transfection, cells were treated with the protein translation inhibitor cycloheximide (CHX) for timepoints of up to 24 hours. In cells expressing control shRNA ∼90% of Mff was degraded within 24 hours, with a half-life of ∼5 hours. In the Parkin knockdown cells, however, the rate of degradation was dramatically slower, increasing the half-life of Mff to ∼24 hours (Fig. 3a,b). Taken together, these data indicate that Parkin-mediated ubiquitination plays a role in physiological Mff degradation under non-stressed conditions.

Parkin ligase activity requires phosphorylation at S65 by PINK1 (22). To further investigate the roles of Parkin and PINK1 in Mff turnover, we generated phospho-null S65A and phospho-mimetic S65D untagged Parkin S65 mutants. We reasoned that Parkin S65A would be unable to efficiently translocate to mitochondria or catalyse ubiquitin-transfer, whereas Parkin S65D would be constitutively active.

HEK293T cells were transfected with WT, S65A or S65D Parkin. Prior to lysis 48 hours post-transfection, cells were treated with CHX or DMSO (vehicle control) for 6 hours. Samples were then Western blotted for Mff (Fig. 3c,d). Consistent with a PINK1-dependent role for Parkin in Mff turnover, expression of Parkin S65D resulted in significant loss of Mff, whereas Parkin S65A had no effect on Mff levels. These data demonstrate that PINK1-mediated phosphorylation and activation of Parkin at S65 is required for its activity in Mff turnover.

### Parkin mediates lysosomal degradation of Mff independent of mitophagy

To establish whether Parkin-mediated degradation of Mff occurs as part of mitophagy we used GFP-Trap® immunoprecipitation to pull down CFP-Mff from cells expressing control or Parkin-targeting shRNA. These samples were then probed for total ubiquitin, K48-linked ubiquitin and K63-linked ubiquitin (Fig. 4,a, b, c, respectively). Indicative of Mff not being a proteasome substrate, no K48-linked ubiquitin was detected on CFP-Mff (Fig. 4b). However, a single K63-linked ubiquitin-reactive species was present (Fig. 4 c), which was reduced in the absence of Parkin, suggesting a role for the lysosome in Parkin-mediated degradation of Mff.

**Fig. 4.**
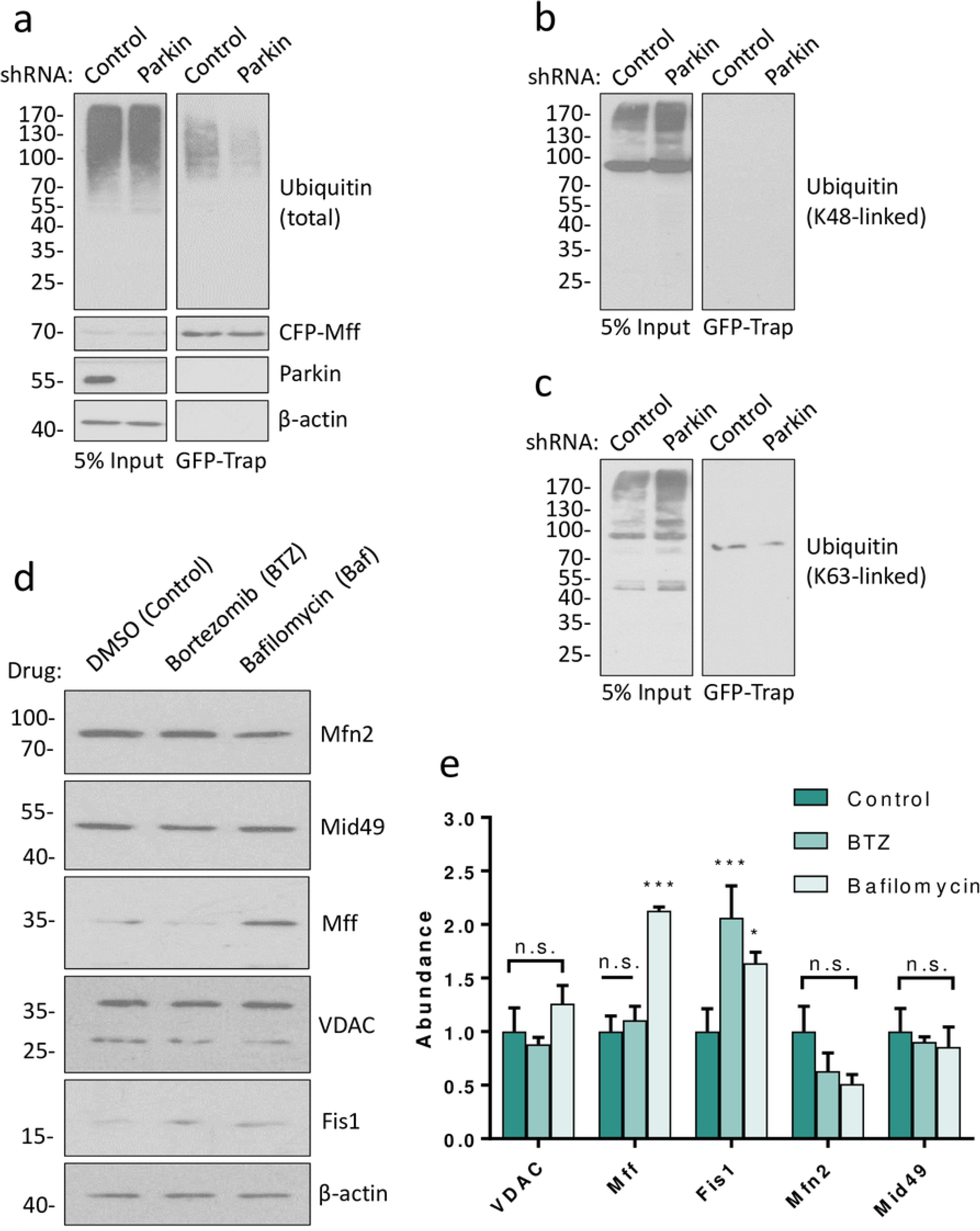
Parkin mediates lysosomal degradation of Mff. a) HEK293T cells were co-transfected with CFP-Mff/GFP and shRNA (scrambled control or Parkin-targeting). Western blots of GFP-immunoprecipitation. As in Figure 2, knockdown of Parkin reduces total Mff ubiquitination. b) Samples from (a), showing no co-immunoprecipitation of K48-linked ubiquitin with CFP-Mff. c) Samples from (a), showing co-immunoprecipitation of a single K63-linked ubiquitinated CFP-Mff species. d) HEK293T cells were transfected with untagged Parkin. Prior to lysis 48 hours post-transfection, cells were treated with DMSO (control), Bortezomib (BTZ, proteasomal inhibitor, 1µM) or Bafilomycin (Baf, lysosomal inhibitor, 100nM) for 6 hours. Lysates were then Western blotted for Mff and other mitochondrial membrane proteins. e) Quantitative analysis of (d), data presented as mean ± SEM. Analysed using ordinary two-way ANOVA with Dunnett’s correction for multiple comparisons. N=3. * p < 0.05, *** p < 0.001.

We next investigated the mechanism of parkin-mediated degradation of Mff by transfecting HEK293T cells with untagged WT Parkin and treated with the proteasomal inhibitor bortezomib (38) or bafilomycin, which inhibits fusion of the autophagosome and lysosome (39). Consistent with Parkin mediating lysosomal degradation of Mff, bafilomycin significantly increased Mff levels, whereas bortezomib treatment had no effect Fig. 4d,e). Interestingly, other MOM proteins tested (Mfn2, Mid49, VDAC) were not affected by bafilomycin, while Fis1 was increased by both proteasomal and lysosomal inhibition (Fig. 4d,e). VDAC and Mfn2 have both been reported to be ubiquitinated by Parkin during mitophagy, yet neither were affected by bortezomib or bafilomycin under the basal conditions of this experiment (32, 40). These data suggest that Parkin-mediated lysosomal degradation of Mff is independent of mitophagy.

## Discussion

Our data show that under basal conditions Parkin ubiquitinates Mff at K151. For this ubiquitination and subsequent and Mff degradation Parkin needs to be activated by PINK1. This Parkin-mediated ubiquitination of Mff coincides with Parkin-mediated Mff degradation, suggesting that Mff turnover is regulated by a Parkin-dependent ubiquitin-mediated pathway. Mff is not a substrate of K48-linked ubiquitination but is a substrate of K63-linked ubiquitination. Furthermore, inhibition of the lysosome, but not the proteasome, recues Mff from Parkin-mediated degradation. These data support a model in which Parkin-mediated degradation of Mff occurs via K63-linked ubiquitination and the lysosome. Interestingly, this activity appears to be distinct from mitophagy, in which depolarised mitochondria recruit Parkin to indiscriminately ubiquitinate MOM proteins prior to their degradation. Of the five MOM proteins assayed in Parkin-overexpressing cells, only Mff and Fis1 were significantly rescued from degradation by inhibition of the lysosome (Fis1 was also rescued by proteasomal inhibition). Mfn2, Mid49 and VDAC were not significantly changed, despite Mfn2 and VDAC being known targets of CCCP-induced, Parkin-mediated degradation, indicating a lack of large-scale mitophagy (32, 40, 41). The degradation of Mff by Parkin under basal conditions, together with its inactivity toward other known substrates under the same conditions, suggest that Parkin-mediated degradation of Mff is a regulatory mechanism independent of mitophagy. Moreover, since PINK1 is maintained at low levels in the MOM of healthy mitochondria, we propose that this mechanism plays a critical background role in maintaining mitochondrial integrity in the absence of induced stress.

Intriguingly, our data demonstrate that K302 is not the sole site of ubiquitination of Mff isoform I and we identified an additional ubiquitination site at K151. However, even in mutants in which both K151 and K302 were ablated residual Mff ubiquitination remained, indicating the presence of at least one other ubiquitination site. Moreover, our observation that ubiquitination of CFP-Mff K151R is unaffected by Parkin knockdown strongly suggests that the sole, or at least predominant, site of Parkin mediated ubiquitination is K151.

Our data, combined with previous reports of Mff ubiquitination and phosphorylation (30, 37) indicate Mff is subject to multiple post-translational modifications. Given its role as the primary receptor for Drp1 in mitochondrial fission, further work will be needed to elucidate how these modifications, and the interplay between them, regulate Mff abundance and activity in health and disease.

## Author contributions

LL devised the project and performed all of the experiments. RS demonstrated MFF K151R has reduced ubiquitination, generated the K302R and 2KR mutants and provided CFP-Mff constructs. KAW supplied new tools, reagents and advice. JMH supervised the project. LL and JMH wrote the manuscript and RS and KAW contributed editing.

## Acknowledgements

We are grateful to the British Heart Foundation (PG/14/60/31014; JMH), Wellcome Trust (studentship to RS), MRC (MR/L003791; KAW, JMH) and BBSRC (BB/R00787X; KAW, JMH) for financial support. The funders had no role in study design, data collection and analysis, decision to publish, or preparation of the manuscript.

